# Iron homeostasis governs erythroid phenotype in Polycythemia Vera

**DOI:** 10.1101/2022.05.03.490556

**Authors:** Cavan Bennett, Victoria E Jackson, Anne Pettikiriarachchi, Thomas Hayman, Ute Schaeper, Gemma Moir-Meyer, Katherine Fielding, Ricardo Ataide, Danielle Clucas, Andrew Baldi, Alexandra L Garnham, Connie SN Li-Wai-Suen, Warren S Alexander, Melanie Bahlo, Kate Burbury, Ashley P Ng, Sant-Rayn Pasricha

## Abstract

Polycythemia Vera (PV) is a myeloproliferative neoplasm driven by activating mutations in *JAK2* that result in unrestrained erythrocyte production, increasing patients’ hematocrit and hemoglobin concentration, placing them at risk of life-threatening thrombotic events. Our GWAS of 440 PV cases and 403,351 controls utilising UK Biobank data found that SNPs in *HFE* known to cause hemochromatosis are highly associated with PV diagnosis, linking iron regulation to PV. Analysis of the FinnGen dataset independently confirmed over-representation of homozygous *HFE* mutations in PV patients. HFE influences expression of hepcidin, the master regulator of systemic iron homeostasis. Through genetic dissection of PV mouse models, we show that the PV erythroid phenotype is directly linked to hepcidin expression: endogenous hepcidin upregulation alleviates erythroid disease whereas hepcidin ablation worsens it. Further, we demonstrate that in PV, hepcidin is not regulated by expanded erythropoiesis but is likely governed by inflammatory cytokines signalling via GP130 coupled receptors. These findings have important implications for understanding the pathophysiology of PV and offer new therapeutic strategies for this disease.

## Introduction

Polycythemia Vera (PV) is a Philadelphia chromosome negative myeloproliferative neoplasm (MPN) driven by activating mutations in *JAK2*^1–4^ that cause unrestrained erythrocyte production, increasing patients’ hematocrit and hemoglobin concentration. Over 95% of cases harbour the JAK2 V617F mutation, with the remainder usually exhibiting a mutation in JAK2 exon 12^5, 6^. Patients usually experience a chronic clinical course^7^, with complications of elevated hematocrit including venous and arterial thrombosis, and systemic symptoms including headache, visual disturbances and pruritis^8^. Therapy for PV typically includes regular venesection to maintain hematocrit below 45%^6, 9^. This phase of disease may continue for years before a proportion of patients develop fibrotic or leukaemic transformation.

Iron availability for erythropoiesis (and other tissues) is governed by hepcidin, the master regulator of systemic iron homeostasis. Hepcidin is produced by the liver and occludes and internalises the sole cellular iron exporter, ferroportin^10, 11^, preventing recycled iron in macrophages and dietary iron absorbed in the intestine from reaching the plasma and hence the erythroid bone marrow^12^. Elevated hepcidin thus reduces iron availability, while suppressed hepcidin enhances it^13^. Systemic iron homeostasis is maintained via transcriptional regulation of hepcidin, whereby iron loading upregulates transcription via the canonical BMP-SMAD signalling pathway^14^. Hepcidin transcription is suppressed by iron deficiency, partly via Matriptase-2 (encoded by *TMPRSS6*) mediated downregulation of Hemojuvelin, a coreceptor for BMP signalling. Increased erythropoiesis also suppresses hepcidin via the erythroid-secreted hormone erythroferrone (ERFE)^15^, which acts to inhibit BMP signalling^16^. Hepcidin is also directly upregulated by inflammation^17^ (via IL6 driven JAK-STAT signalling^18^).

Systemic iron metabolism and PV may be closely intertwined.^19^ Over 50% of PV patients present with iron deficiency at diagnosis^20^, potentially limiting or even masking the disease. Venesection induces iron deficiency to limit further erythropoiesis. Recent reports have indicated that treatment of PV patients with hepcidin analogues^21^ or pharmaceutical upregulation of hepcidin in pre-clinical models of PV^22, 23^ may ameliorate disease phenotype. Given the emerging interest in manipulation of iron homeostasis in PV, comprehensive characterisation of the regulation of hepcidin and its role in PV disease is imperative.

Here, we establish an association between disordered systemic iron homeostasis and risk of PV diagnosis through genome-wide association analyses. Using multiple preclinical models of JAK2 V617F driven PV, we then show that hepcidin levels are critical to governing the severity of the erythroid phenotype. Further, we demonstrate that in PV, hepcidin is not regulated by erythropoiesis but likely governed by inflammatory cytokines signalling through GP130 couple receptors. These findings provide novel insights into understanding the pathophysiology of PV and have important implications for new therapeutic interventions.

## Methods

Full details are available in the Supplemental Materials.

### Ethics

Patient samples were collected under Walter and Eliza Hall Institute (WEHI) ethics 18/10LR; all participants provided written informed consent as per the Declaration of Helsinki. The UK Biobank has approval from the UK North West Multi-centre Research Ethics Committee (MREC) as a Research Tissue Bank (approval 21/NW/0157). The use of mice was in accordance with requirements set out by WEHI Animal Ethics Committee (approvals 2017.031 and 2020.034).

### UK Biobank GWAS

We undertook a genome-wide association study (GWAS) of PV cases versus controls. Associations with single nucleotide polymorphisms (SNPs) and small indels were tested genome-wide, using Regenie^24^, under an additive genetic model. Associations included adjustment for sex, age, genotyping array, 10 ancestry principal components, and relatedness. GWAS results were filtered to include only variants with a minor allele frequency (MAF) >= 1.2%. For variants within the *HFE* locus, associations with PV were also tested assuming a recessive model, with covariate adjustment as above. Associations at this locus and the four blood cell traits were tested separately in PV cases and controls, using Regenie, as above, under both the additive and recessive models.

### FinnGen GWAS analysis

We utilised the FinnGen resource, data release 6^25^. PV cases were identified as having a relevant ICD-8, 9 or 10 code in the hospital discharge register, cause of death register, or cancer register; controls were non-PV individuals without a record of cancer. Genome-wide associations with PV were carried out with adjustment for sex, age, 10 ancestry PCs and genotyping batch, and assuming an additive genetic model. The difference in the proportion of homozygous AA genotype individuals in PV cases vs controls was tested using Fisher’s Exact Test.

### Animals

Erythroferrone knockout (*Erfe*-KO)^16^ and inducible hepcidin knockout (iHamp-KO)^26^ mice have been described previously. Transgenic mice with a single copy Cre recombinase-dependent *Jak2*-V617F transgene located downstream of the *Col1a1* locus (LSL-Jak2-V617F; CreERT2^T/+^) were generated using the strategy developed by Beard et al ^27^ (full methodology in Supplemental Materials). Age, sex matched control animals were used in all experiments.

### Bone marrow transplant model of Polycythemia Vera

LSL-Jak2-V617F; CreERT2^T/+^ (PV) or LSL-Jak2-V617F lacking CreERT2 (control) bone marrow (BM) cells were intravenously injected into lethally irradiated Ly5.1/J (B6.SJL-Ptprca Pepcb/BoyJ) recipient mice. Seven weeks post BM transplantation, mice were given tamoxifen (Sigma; 4.2mg in 90% corn oil/10% ethanol) by oral gavage on 2 consecutive days to induce expression of the mutant *Jak2* allele. Full details in Supplemental Materials.

### Administration of drugs, antibodies, and siRNA

*TMPRSS6* siRNA (Silence Therapeutics GmbH, Berlin, Germany) comprised a double-stranded 19mer RNA oligonucleotide targeting human *TMPRSS6,* linked to a GalNAc unit at the 5′ end of the sense strand enabling hepatic targeting^28^. *TMPRSS6* siRNA was diluted in sterile PBS and 5mg/kg (or an equal volume of PBS) administered by subcutaneous injection every 3 weeks for a total of 3 doses.

500μg anti-mouse IL6 (clone MPF-20F3) or Rat IgG1 Kappa control antibodies (both made in house) were administered by intraperitoneal injection every 3 days for a total of 7 doses.

### RNA-Seq

300ng RNA was used as input for indexed libraries according to Illumina’s TruSeq RNA Sample Prep guidelines. The indexed libraries were pooled and sequenced (paired end, 2x 76 cycles) on a NextSeq 500 instrument. Paired-end RNA sequencing reads were aligned to the mm10 build of the mouse reference genome using Rsubread package v2.4.3^29^. Differential expression analyses were carried out using limma v3.48.3^30, 31^ and edgeR v3.34.0^32^. False discovery rate was controlled below 5% using the Benjamini and Hochberg method. The mean-difference plot was generated using the limma’s plotMD function while the heatmap was created using the pheatmap software package.

Pathway analyses were performed on differentially expressed genes to test for overrepresentation of biological pathways as defined by Gene Ontology (GO)^33, 34^ and Kyoto Encyclopedia of Genes and Genomes (KEGG) pathways^35–37^ using limma’s goana and kegga functions respectively. Analysis of the Molecular Signatures Database (MSigDB) hallmark gene sets^38, 39^ was undertaken using the fry gene set test in limma. See Supplemental Materials for full methodology.

### Statistical analysis

Sample sizes and statistical tests for each experiment are denoted in the figure legends. Statistical testing was performed in Prism 9.3.1; GraphPad Software.

For original data contact the corresponding authors. Additional methods are in the Supplemental Material.

## Results

### Genome-wide association analysis links PV diagnosis with HFE variants

To investigate potential genetic associations in PV patients with molecular regulators of iron metabolism, we undertook a GWAS of 440 PV cases and 403,351 controls in the UK Biobank (fig. 1A). We tested 9,191,064 variants genome-wide (fig. 1B). Three genetic loci had SNPs with genome-wide significant (p<5E-8) associations. Table 1A details the top SNP at each locus. The most significant association was with rs62541556 (p=8.85E-21), a germline SNP in *JAK2* that tags the 46/1 haplotype implicated in MPNs and is associated with increased susceptibility to V617F-mutant clonal hematopoiesis^40, 41^. The other genome-wide significant associations were with rs79220007 in *HFE* (p=1.42E-14) and rs3836364 in *FBLN2* (p=4.75E-8).

**Figure 1:**
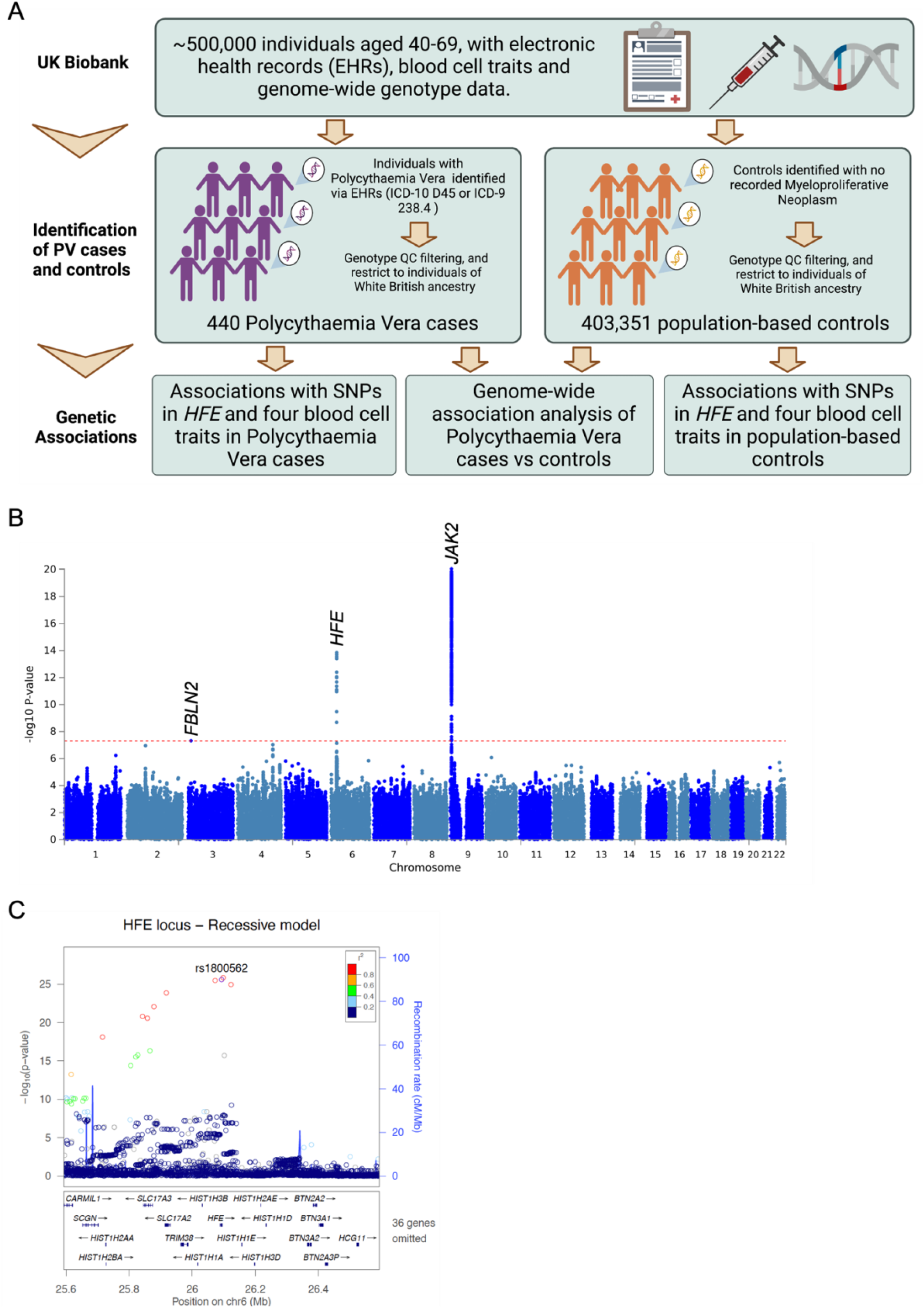
Genome-wide association study (GWAS) of Polycythemia Vera. (A) Schematic of GWAS. Created with BioRender.com. (B) Manhattan plot showing results of GWAS of 440 PV cases vs 403,351 controls, assuming an additive genetic model. Red dotted line show genome-wide significance level (p<5E-8). Three loci with associations exceeding this threshold are labelled with the nearest gene. (C) LocusZoom plot of associations at the *HFE* locus, assuming a recessive genetic model. rs1800562 (C282Y) highlighted, with the other single nucleotide polymorphisms (SNPs) coloured according to linkage disequilibrium (r^2^) to that SNP.

**Table 1.** A: Genome-wide significant loci from a GWAS of PV, assuming an additive genetic model

| Gene | ID | Chr:Pos | Ref allele /<br>Effect allele | Effect allele<br>Frequency | Odds Ratio (SE<sub>beta</sub>) | P |
| --- | --- | --- | --- | --- | --- | --- |
| *JAK2* | rs62541556 | 9:5049092 | G/T | 25.6% | 1.870 (0.067) | 8.85E-21 |
| *HFE* | rs79220007 | 6:26098474 | T/C | 7.7% | 2.128 (0.098) | 1.42E-14 |
| *FBLN2* | rs3836364 | 3:13676875 | TAC/T | 25.1% | 1.505 (0.075) | 4.746E-8 |

5 B: Association of rs1800562 (C282Y) variant in *HFE* with PV, under the additive, and recessive genetic models
| ID | Chr:Pos | Ref allele /<br>Effect allele | Genotype<br>distribution cases | Genotype<br>distribution<br>controls | Additive<br>model Odds<br>Ratio (SE<sub>beta</sub>) | Additive<br>model P | Recessive<br>model Odds<br>Ratio (SE<sub>beta</sub>) | Recessive model<br>P |
| --- | --- | --- | --- | --- | --- | --- | --- | --- |
| **rs1800562** | 6:26093141 | G/A | 340 / 70 / 35 | 246,662 /<br>58,305 / 2596 | 2.106 (0.098) | 2.91E-14 | 12.581 (0.239) | 2.632E-26 |

Chr:Pos – chromosome and position based on hg19
Odds ratio – in respect of effect allele
SE<sub>beta</sub> – standard error of the beta (log odds ratio).
Genotype distribution: counts of individuals with G/G homozygous, G/A heterozygous and A/A homozygous genotypes, in that order.

Interestingly, the top SNP in *HFE*, rs79220007, is in very high linkage disequilibrium (r^2^=1.0) with rs1800562, a SNP known to cause the C282Y mutation, the most common mutation causing Type I Hemochromatosis, an autosomal recessive disorder producing iron overload^42^. Amongst PV cases, there was an excess of individuals with the AA homozygous genotype, compared to what would be expected under Hardy-Weinberg equilibrium (2.7 expected vs 35 observed). Given the over-representation of AA genotypes in PV cases, and the established role of homozygous C282Y mutations in disease, we re-examined the *HFE* locus for associations with PV under a recessive model. The association with rs1800562 was more statistically significant under the recessive model than the additive model (p=2.63E-26 vs p=2.91E-14; table 1B, fig. 1C) and the homozygous AA genotype was associated with 12.58 times increased odds of PV, compared to G/G or G/A genotypes.

We next sought further evidence for the association of rs1800562 with PV through a look-up of rs1800562 in a GWAS of PV in the FinnGen study (394 PV cases vs 217,902 controls)^25^. Again, there was an over-representation of the AA genotype amongst PV cases (0.6 expected vs 4 observed). The crude odds ratio comparing the frequency of the AA homozygous genotype in PV cases vs controls in FinnGen was 5.19 (p=8.29E-3, Supplemental Table 1). The FinnGen GWAS assumed an additive genetic model and did not demonstrate an association between rs1800562 and PV (p=0.68). However, the C282Y variant is less frequent in Finnish compared to British populations (MAF 3.7% vs 7.8%), therefore, there may have limited power to capture the recessive association where an additive effect is assumed.

Using the UK Biobank, we then examined associations between rs1800562 and blood cell traits in PV cases and control individuals separately under both additive and recessive models. In line with a previous GWAS^43^, in control individuals the A allele of rs1800562 was highly associated with higher hemoglobin concentrations, hematocrits and mean corpuscular volume, but lower erythrocyte count (Supplemental Table 2); this latter association was larger and more statistically significant under a recessive model (Supplemental Table 2).

HFE is involved in the regulation of iron homeostasis through influencing regulation of hepcidin transcription (reviewed^44^). The C282Y HFE mutation dysregulates the iron-hepcidin axis, causing lower hepcidin concentrations relative to iron stores^45^. We therefore sought to further investigate how iron homeostasis may affect PV disease severity by studying hepcidin regulation in a mouse model of PV.

### Hepcidin is not suppressed in murine Polycythemia Vera

To define hepcidin regulation in PV we developed a BM transplant model of PV (fig. 2A). Importantly, this model allows mutant *Jak2*-V617F expression in the hematopoietic lineage whilst preserving canonical wildtype JAK2-STAT signalling in hepatocytes. Reconstitution of the BM by donor cells was highly efficient (Supplemental Figure 1). Ten weeks after tamoxifen induction, recipient mice transplanted with LSL-Jak2-V617F; CreERT2^T/+^ BM (hereafter, PV mice) exhibited a classic PV phenotype with increased red cells, hemoglobin, hematocrit, decreased mean cell volume (MCV), lack of erythropoietin (EPO) expression, and splenomegaly (fig. 2B-G). In addition, compared to controls PV mice had a skewed distribution of BM resident erythropoietic progenitor cells, with increased levels of the most mature (stage V) erythroid cells (fig. 2H). Interestingly, despite an increase in erythropoiesis and higher *Erfe* mRNA and protein levels (fig. 2I-J), hepcidin mRNA (gene: *Hamp1*) and protein levels were not reduced in the PV mice (fig. 2K-L). There was no difference in expression of other BMP-SMAD target genes, namely *Atoh8*, *Smad7* and *Id2* (fig. 2M-O).

**Figure 2:**
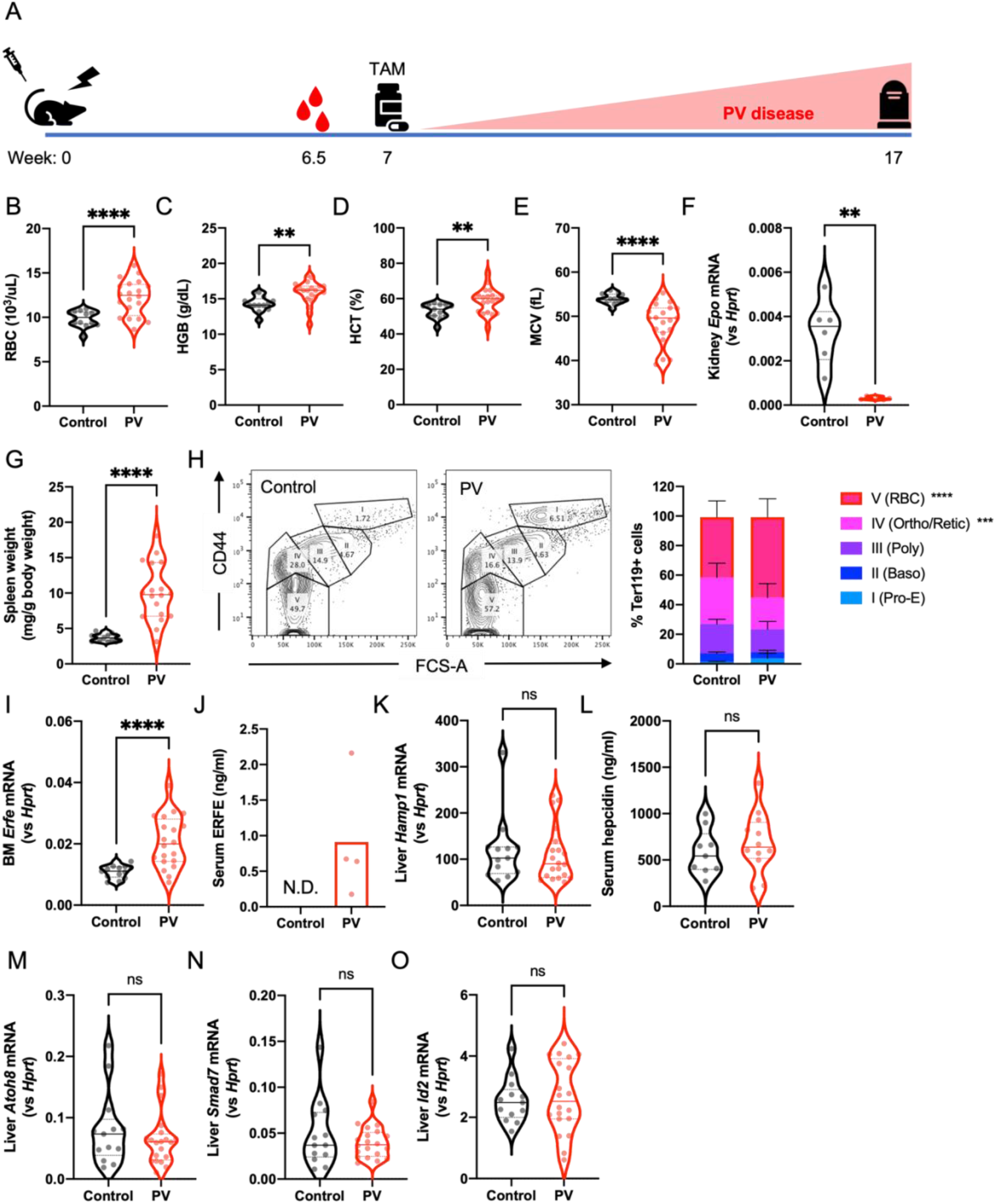
Hepcidin is not suppressed in a novel mouse model of Polycythemia Vera. (A) Schematic of BMT PV mouse model. At week 0, recipient mice were lethally irradiated and injected with donor bone marrow cells. Recipient mice were bled from the retro-orbital plexus at 6.5 weeks to determine chimerism. At 7 weeks mice were treated with tamoxifen (TAM) to induce *Jak2*-V617F expression. PV disease phenotype increases over 10 weeks with mice being humanely euthanised at week 17. (B-E) Red blood cells (RBC; B), hemoglobin (HGB; C), hematocrit (HCT; D) and mean cell volume (MCV; E) determined by automated hemocytometer on EDTA anticoagulated whole blood. N=13 control/20 PV. (F) Kidney *Epo* mRNA expression relative to *Hprt* determined by RT-qPCR. N=6. (G) Spleen weight normalised to total body weight. N=13 control/18 PV. (H) Representative flow cytometry plots showing CD44 expression against Forward/Side Scatter Area (FSC-A) of Ter119 expressing (Ter119+) bone marrow cells. Based on CD44 expression and FSC-A, cells were gated into 5 distinct populations: I – proerythroblast (Pro-E), II – basophilic erythroblasts (Baso), III – polychromatic erythroblasts (Poly), IV – orthochromatic erythroblasts and reticulocytes (Ortho/Retic), and V – red blood cells (RBC). N=13 control/20 PV. (I) Bone marrow (BM) *Erfe* mRNA expression relative to *Hprt* determined by RT-qPCR. N=13 control/20 PV. (J) Serum ERFE determined by enzyme-linked immunosorbent assay. N=5 control/4 PV. (K) Liver *Hamp1* mRNA expression relative to *Hprt* determined by RT-qPCR. N=13 control/20 PV. (L) Serum hepcidin determined by enzyme-linked immunosorbent assay. N=9 control/12 PV. (M-O) Liver *Atoh8* (M), *Smad7* (N) and *Id2* (O) mRNA expression relative to *Hprt* determined by RT-qPCR. N=13 control/20 PV. Unpaired 2-tailed t-test with Welch’s correction (B-G, I, L and O), Mann-Whitney test (K, M and N) or two-way ANOVA with Šídák’s correction for multiple comparisons (H). ** p>0.01; ***p>0.001; ****p>0.0001; ns = non-significant; N.D. = not detected.

### ERFE does not regulate hepcidin expression in PV

ERFE is produced by erythroblasts^15^ and acts as a BMP6 ligand trap^16^, decreasing BMP-SMAD signalling and thus hepatic hepcidin expression. Since PV causes increased erythropoiesis, we confirmed increased ERFE levels in our PV model (fig. 2J): a finding we also observed in a cohort of PV patients who demonstrated raised ERFE levels compared to healthy controls (fig. 3A). To determine the impact of ERFE on both hepcidin levels and disease severity in PV we crossbred the LSL-Jak2-V617F; CreERT2^T/+^ mice with a previously published *Erfe* knockout (*Erfe*-KO) mouse ^16^ (PV x *Erfe*-KO mice). The inter-crossed mice were used as donors for BM transplants allowing *Jak2*-V617F expression along with ablated *Erfe* in hematopoietic cells of recipient mice. While PV mice have increased ERFE levels (fig. 2J), ERFE production is completely ablated in PV x *Erfe*-KO mice (fig. 3B). Deletion of *Erfe* in PV mice did not alter *Hamp1* expression (fig. 3C), hepcidin protein levels (fig. 3D), or change expression of other BMP-SMAD target genes (fig. 3E). Deletion of *Erfe* in PV mice did not alter severity of PV disease, with no change in red cells, hemoglobin, or hematocrit (fig. 3F-H).

**Figure 3:**
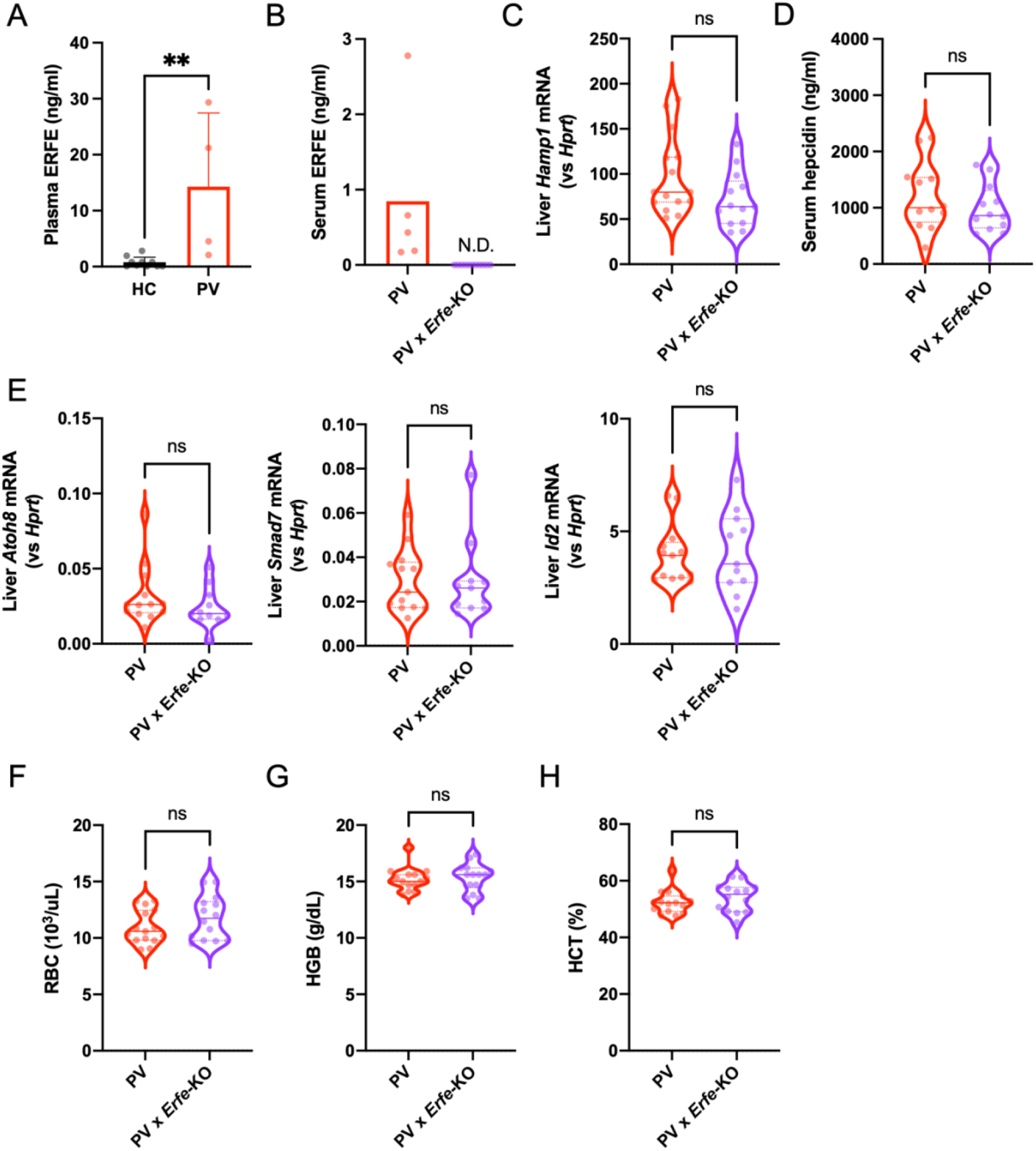
ERFE does not affect hepcidin in Polycythemia Vera. (A) Plasma ERFE of PV patient and healthy control (HC) donors determined by enzyme-linked immunosorbent assay. N=10 HC/4 PV. (B) Mouse serum ERFE determined by enzyme-linked immunosorbent assay. N=5 PV/11 PV x Erfe-KO. (C) Liver *Hamp1* mRNA expression relative to *Hprt* determined by RT-qPCR. N=15 PV/13 PV x Erfe-KO. (D) Mouse serum hepcidin determined by enzyme-linked immunosorbent assay. N=12 PV/12 PV x Erfe-KO. (E) Liver *Atoh8* (left panel), *Smad7* (middle panel), and *Id2* (right panel) mRNA expression relative to *Hprt* determined by RT-qPCR. N=13 PV/11 PV x Erfe-KO. (F-H) Red blood cells (RBC; F), hemoglobin (HGB; G), and hematocrit (HCT; H) determined by automated hemocytometer on EDTA anticoagulated whole blood. N=15 PV/14 PV x Erfe-KO. Mann-Whitney test (A, C, E) or Unpaired 2-tailed t-test with Welch’s correction (B, D, F-H). ** p>0.01; ns = non-significant; HC = healthy control; N.D. = not detected.

### Hepcidin determines PV erythroid disease severity

We hypothesised that hepcidin regulates erythroid disease in PV. To test this hypothesis, we first deleted hepcidin in PV mice, using a previously published inducible hepcidin knockout (iHamp-KO) mouse model^26^. Since mutant JAK2-V617F expression in hepatocytes was not a concern in mice unable to generate hepcidin, we crossbred LSL-Jak2-V617F; CreERT2^T/+^ mice with iHamp-KO mice, creating mice with simultaneously inducible systemic mutant *Jak2*-V617F expression and hepcidin deletion (PV x iHamp-KO). As expected, PV x iHamp-KO mice showed no *Hamp1* expression (fig. 4A) and greatly reduced hepcidin protein (fig. 4B). Importantly, the PV x iHamp-KO mice developed a more severe PV erythroid phenotype than their hepcidin expressing PV littermates, as manifested by increased hemoglobin concentration (19.20 vs 22.70 g/dL, p=0.0021; fig. 4C) and hematocrit (68.70 vs 75.85%, p=0.0065 fig. 4D). This may reflect increased iron availability for red cell production as evidenced by increased MCV (43.98 vs 57.62 fl, p=0.0012; fig. 4E). PV x iHamp-KO mice had fewer erythrocytes than their hepcidin expressing counterparts (15.78 vs 13.16×10^3^/μL, p=0.0052; fig 4F), which, interestingly, recapitulates blood cell traits in control individuals with the *HFE* rs1800562 A allele (Supplemental Table 2).

**Figure 4:**
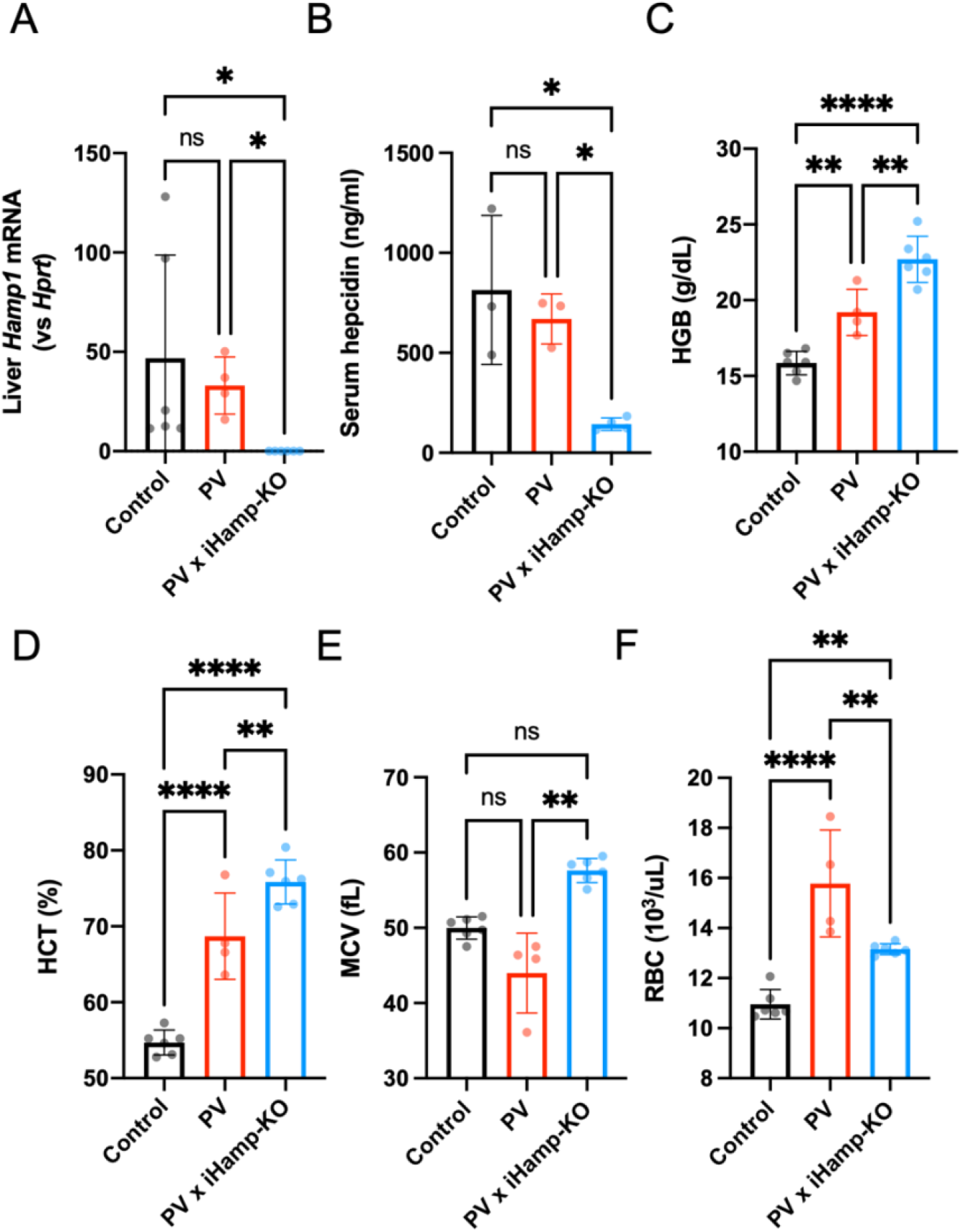
Hepcidin deletion worsens PV erythroid disease severity. (A) Liver *Hamp1* mRNA expression relative to *Hprt* determined by RT-qPCR. N=6 control and PV /4 PV x iHamp-KO. (B) Mouse serum hepcidin determined by enzyme-linked immunosorbent assay. N=3 control and PV x iHamp-KO/4 PV. (C – E) Hemoglobin (HGB; C), hematocrit (HCT; D), mean cell volume (MCV; E) and red blood cells (RBC; F) determined by automated hemocytometer on EDTA anticoagulated whole blood. N=6 control and PV x iHamp-KO /4 PV. Kruskal-Wallis test with Dunn’s correction for multiple comparisons (A and E) or Ordinary one-way ANOVA with Holm-Šídák’s correction for multiple comparisons (B-D and F). *p>0.05; ** p>0.01; ****p>0.0001; ns = non-significant.

### TMPRSS6 inhibition increases endogenous hepcidin and improves PV disease

Since hepcidin ablation worsened the PV erythroid phenotype, we hypothesised that increasing hepcidin levels would impair erythroid iron availability and reduce erythroid disease. Injectable hepcidin mimetics (e.g. LJPC-401 and PTG-300) have a relatively short therapeutic duration (<7 days). We sought to raise endogenous hepcidin expression over a prolonged period. We hypothesised that downregulation of *Tmprss6* (a negative regulator of hepcidin expression^46^) would increase hepcidin expression in our PV model. Mice were injected with *TMPRSS6* siRNA (that has been shown to cause sustained downregulation of murine *Tmprss6*^28^) or PBS every 3 weeks for a total of 3 injections (fig. 5A). *TMPRSS6* siRNA treatment resulted in efficient knockdown of hepatic *Tmprss6* compared to animals treated with PBS (fig. 5B). Decreased *Tmprss6* expression led to a 2.02-fold increase in hepatic *Hamp1* (fig. 5C) and a 2.47-fold increase in hepcidin protein (fig. 5D) in PV mice treated with *TMPRSS6* siRNA compared to PV mice treated with PBS.

**Figure 5:**
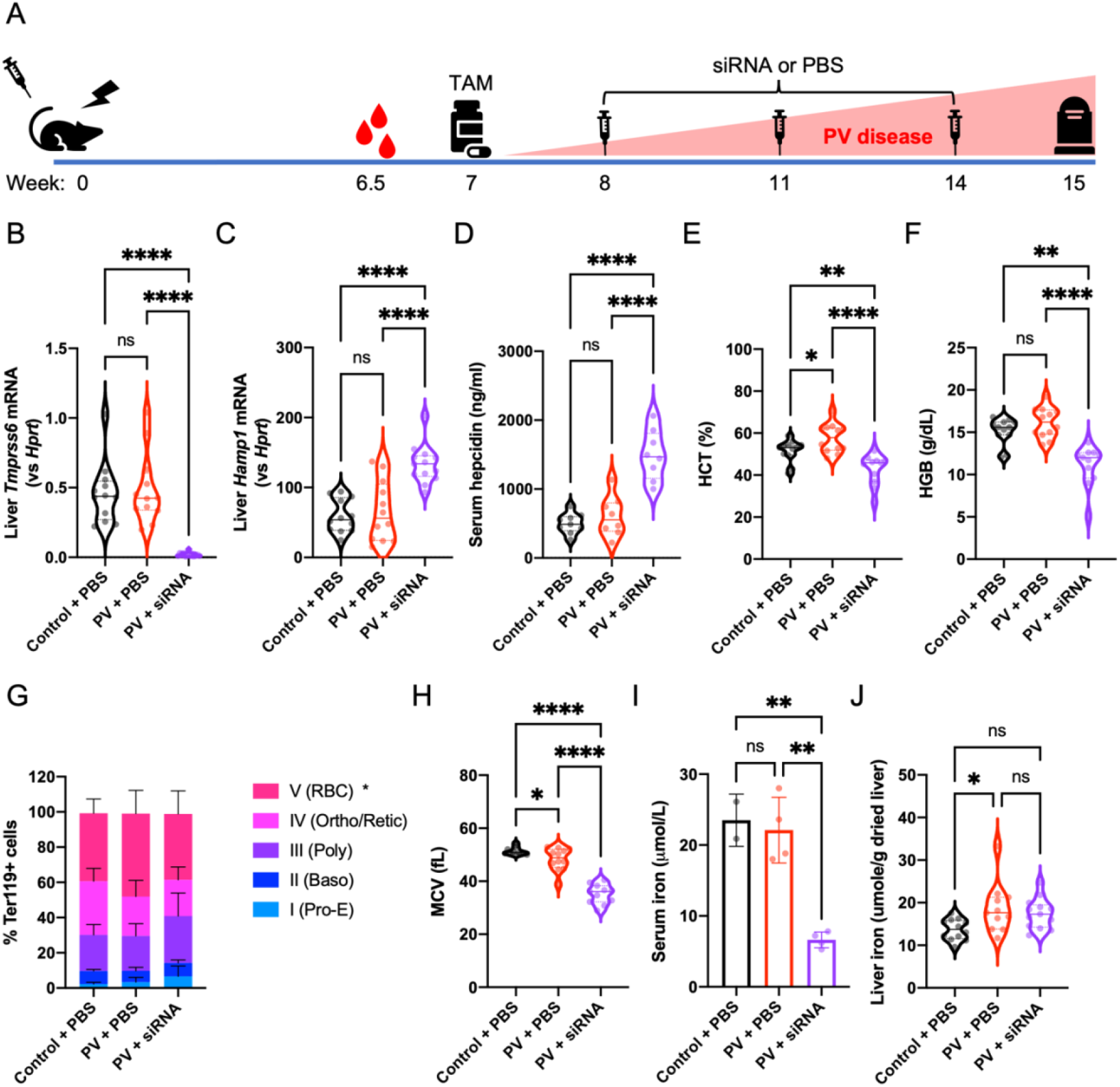
*TMPRSS6* inhibition increases endogenous hepcidin and improves PV disease severity. (A) Schematic of experimental design. At week 0, recipient mice were lethally irradiated and injected with 2.5×10^6^ donor cells. Recipient mice were bled from the retro-orbital plexus at 6.5 weeks to determine chimerism. At 7 weeks mice were treated with tamoxifen (TAM) to induce *Jak2*-V617F expression. Mice were treated with *TMPRSS6* siRNA or PBS at weeks 8, 11 and 14. PV disease phenotype increases over 8 weeks with mice humanely euthanised at week 15. (B – C) Liver *Tmprss6* (B) and *Hamp1* (C) mRNA expression relative to *Hprt* determined by RT-qPCR. N=12 PBS groups/13 siRNA group. (D) Mouse serum hepcidin determined by enzyme-linked immunosorbent assay. N=9 control group/8 PV groups. (E – F) Hematocrit (HCT; E) and hemoglobin (HGB; F) determined by automated hemocytometer on EDTA anticoagulated whole blood. N=12 PBS groups/11 siRNA group. (G) Bone marrow cells flushed from the tibia and femur were incubated with antibodies against CD44 and Ter119. Ter119 positive cells were gated into 5 distinct populations based on CD44 expression and Forward/Side Scatter Area: I – proerythroblast (Pro-E), II – basophilic erythroblasts (Baso), III – polychromatic erythroblasts (Poly), IV – orthochromatic erythroblasts and reticulocytes (Ortho/Retic), and V – red blood cells (RBC). N=12 PBS groups/13 siRNA group. (H) Mean cell volume (MCV) determined by automated hemocytometer on EDTA anticoagulated whole blood. N=12 PBS groups/11 siRNA group. (I) Serum iron. N=2 control group / 4 PV groups (J) Liver non-heme iron. N=12 control/10 PV + PBS/13 PV + siRNA. Kruskal-Wallis test with Dunn’s correction for multiple comparisons (B, F) or Ordinary one-way ANOVA with Holm-Šídák’s correction for multiple comparisons (C-E, H-J). For (G) PV + PBS group was compared to PV + siRNA group only by two-way ANOVA with Šídák’s correction for multiple comparisons. *p>0.05; ** p>0.01; ****p>0.0001; ns = non-significant.

Hepcidin upregulation by *TMPRSS6* siRNA treatment significantly decreased hematocrit (57.58 vs 42.68%, p<0.0001; fig. 5E) and hemoglobin (16.13 vs 11.07 g/dL, p<0.0001; fig. 5F) in PV mice. *TMPRSS6* siRNA treatment normalised the distribution of BM resident erythropoietic progenitor cells, reducing the mature stage V population (fig. 5G). MCV was also reduced in PV mice treated with *TMPRSS6* siRNA (fig. 5H) indicative of iron-restricted erythropoiesis. Indeed, *TMPRSS6* siRNA treatment significantly reduced serum iron in PV mice (fig. 5I). However, this treatment did not reduce liver iron (fig. 5J).

### Inflammatory cytokines may upregulate hepcidin in PV

PV is considered a disease of chronic inflammation, with several inflammatory cytokines elevated in PV^47, 48^. Above, we showed that BMP-SMAD signalling is unchanged in our PV model. However, hepcidin transcription is also regulated by JAK-STAT signalling, mediated through inflammatory pathways, classically via IL6^18^. Thus, we examined whether JAK-STAT signalling was increased in hepatocytes of PV mice. RNA-Seq analysis on livers from BM transplanted PV and control animals revealed 1466 (598 downregulated; 868 upregulated) differentially expressed genes (DEGs; fig. 6A and Supplemental Table 3) in PV mice compared to controls. The transcriptional profile of livers from PV mice was different to that of control animals (fig. 6B). Gene Ontology (GO) pathway analysis of DEGs revealed upregulation of biological processes related to JAK-STAT signalling in PV mice compared to control (fig. 6C, top 3 bars). Consistent with this analysis, gene set enrichment analysis (GSEA) using the MSigDB hallmark gene sets revealed upregulation of the inflammatory response pathway in the liver of PV mice (fig. 6C, fourth bar).

**Figure 6:**
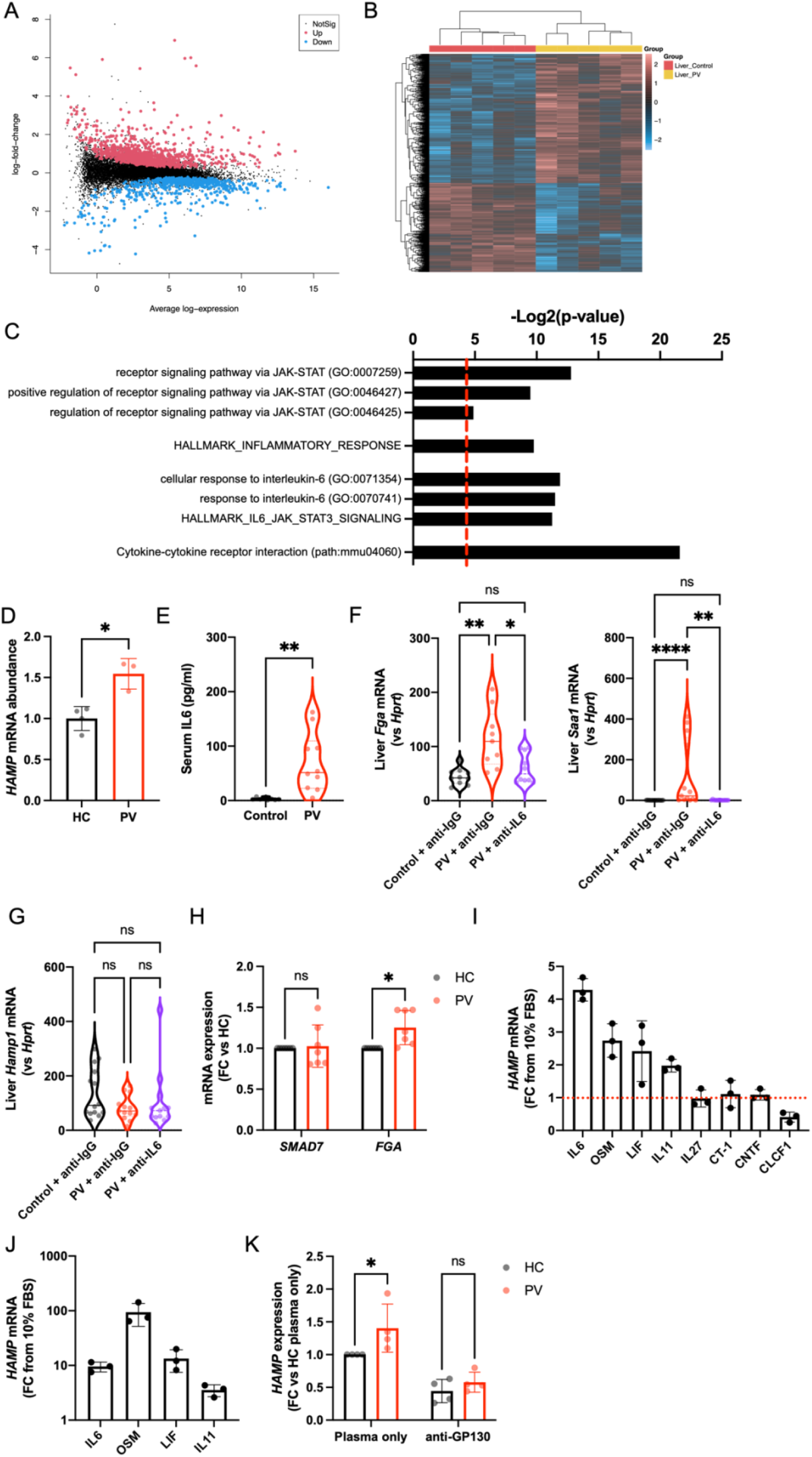
Inflammatory cytokines may upregulate hepcidin in Polycythemia Vera. (A) Mean-difference plot showing the average log-expression of each gene (x-axis) and their log-fold change between PV and control liver samples (y-axis). The differentially expressed genes (DEGs) are highlighted with points in red and blue indicating upregulated and downregulated genes, respectively (adjusted p-value <0.05). (B) Heatmap of the expression of all DEGs with hierarchical clustering where expression values are standardised to have mean 0 and standard deviation 1 for each gene. (C) Bar chart depicting Gene Ontology (GO) biological processes, MSigDB hallmark gene sets or KEGG pathways relating to JAK-STAT signalling, inflammatory response, interleukin-6 (IL6) responses or cytokine-cytokine receptor interactions that are associated with upregulated genes in PV liver samples vs control. X-axis represents statistical significance of the enrichment, increasing from left to right. Red dashed line represents p=0.05. (D) *HAMP* mRNA expression determined by RT-qPCR of HepG2 cells cultured in media supplemented with 2% plasma from healthy control donors (HC) or PV patients. N=4 HC/3 PV. (E) Mouse serum interleukin-6 (IL6) determined by enzyme-linked immunosorbent assay. N=8 Control/10 PV. (F) Liver *Fga* (left panel) and *Saa1* (right panel) relative to *Hprt* determined by RT-qPCR. N=10 Control + anti-IgG and PV +anti-IL6/9 PV + anti-IgG. (G) Liver *Hamp1* relative to *Hprt* determined by RT-qPCR. N=14 Control group/13 PV groups. (H) *SMAD7* and *FGA* mRNA expression determined by RT-qPCR of HepG2 cells cultured in media supplemented with 2% plasma from healthy control donors (HC) or PV patients. N=7. (G-H) *HAMP* mRNA expression determined by RT-qPCR of HepG2 (G) or Huh7 (H) cells cultured in media supplemented with 10ng/ml recombinant human interleukin-6 family cytokines. N=3. The red line in (G) indicates *HAMP* expression in the absence of additional cytokines. (I) *HAMP* mRNA expression determined by RT-qPCR of HepG2 cells cultured in media supplemented with 2% plasma from healthy control donors (HC) or PV patients with or without the addition of anti-GP130 antibodies. N=4. IL6 – Interleukin-6; OSM – Oncostatin M; LIF – Leukemia inhibitory factor, IL-11 – Interleukin-11; IL-27 – Interleukin-27, CT-1 – Cardiotrophin 1; CNTF – Ciliary neurotrophic factor, CLCF1 – Cardiotrophin-like cytokine factor 1. Unpaired 2-tailed t-test with Welch’s correction (D, E), Kruskal-Wallis test with Dunn’s correction for multiple comparisons (F, G) or Two-way ANOVA with Šídák’s correction for multiple comparisons (H, K). *p>0.05; ** p>0.01; ****p>0.0001; ns = non-significant.

We thus explored the hypothesis that soluble factors (including cytokines) raise hepcidin levels in PV patients using a human *in vitro* hepatocyte model. Liver-derived HepG2 cells grown in media supplemented with plasma from PV patients exhibited increased hepcidin (*HAMP*) mRNA expression compared to cells grown in media supplemented with healthy donor plasma (fig. 6D), suggesting that PV patients produce soluble factors that upregulate hepcidin. We reasoned that IL6 may be driving this upregulation, since PV mice exhibited increased serum IL6 (fig. 6E). In addition, RNA-seq GO pathway analysis, as well as GSEA using the MSigDB hallmark gene sets, revealed increased IL6 driven responses in the livers of PV mice (fig. 6C, fifth to seventh bars). To determine whether IL6 alone is responsible for hepcidin upregulation in PV, BM transplanted mice were treated with anti-IL6 or control (anti-IgG) antibodies. Mice treated with anti-IL6 antibodies showed normalisation of hepatic JAK-STAT (e.g. *Fga*) and IL6-JAK-STAT (e.g. *Saa1*) transcripts (fig. 6F), confirming IL6 signalling neutralisation. However, IL6 neutralisation did not alter hepcidin expression (fig. 6G).

We therefore hypothesised that other cytokines may upregulate hepcidin in PV. In keeping with this, KEGG pathway analysis of DEGs revealed upregulation of genes involved in cytokine-cytokine receptor interaction in the liver of PV mice compared to controls (fig. 6C, bottom bar). HepG2 cells cultured in media supplemented with plasma from PV patients exhibited increased JAK-STAT target gene expression (*FGA*) but no change in BMP-SMAD target gene expression (*SMAD7*) when compared to cells cultured in media supplemented with plasma from healthy donors (fig. 6H). Notably, levels of some IL6-family cytokines (IL11^49, 50^, OSM^51^) are known to be elevated in PV. Thus, we explored a role for other IL6-family cytokines in hepcidin regulation since these cytokines induce JAK-STAT signalling via a common GP130 receptor subunit (except IL31). Addition of individual IL6-family cytokines to HepG2 cells revealed that in addition to IL6, IL11, OSM and LIF increase *HAMP* mRNA levels; whereas IL27, CT-1 and CNTF have no effect and CLCF1 decreases hepcidin expression (fig. 6I). The increase in hepcidin expression by IL11, OSM and LIF (as well as IL6) was confirmed in a second hepatocyte cell line, Huh7 (fig. 6J). We then determined whether inhibition of GP130 could normalise hepcidin expression in HepG2 cells treated with PV plasma. HepG2 cells grown in media supplemented with PV plasma and anti-GP130 antibodies no longer exhibited increased *HAMP* expression, rather *HAMP* expression decreased, reaching levels indistinguishable to that of cells cultured in media supplemented with control plasma and anti-GP130 antibodies (fig. 6K).

## Discussion

Polycythemia Vera is a chronic blood cancer driven by activating JAK2 mutations that cause unrestrained erythrocyte production. Here, we provide population genetic and experimental evidence that diagnosis and clinical features of PV are influenced by systemic iron homeostasis. Taken together, this work establishes a central role of iron regulation in influencing the erythroid phenotype in PV and provides a clinical rationale for the use of therapies that modify iron metabolism in the treatment of this disease.

Our GWAS implicates the *HFE* locus as a key region associated with PV diagnosis, formally linking systemic iron regulation and PV. The top SNP in *HFE* associated with PV, rs7922007, is in very high linkage disequilibrium with rs1800562, the pathogenic HFE C282Y mutation (which causes recessive hemochromatosis) and is the likely driver of our observed association. Interestingly, this disease-causing SNP is not associated with the diagnosis of the other MPNs (Supplemental Table 4), indicating iron metabolism is linked to PV exclusively. These HFE mutations disrupt the iron-hepcidin axis, resulting in excess iron absorption^45, 52^. Treatment of PV with iron may accelerate increases in hematocrit, thus worsening disease^53^. Conversely, iron deficiency is common at diagnosis in patients with PV^20^, and may conceal diagnosis^54^. We hypothesise that PV patients with *HFE* mutations have relatively suppressed hepcidin, which supports iron availability for erythropoiesis, increasing the likelihood of clinical presentation and diagnosis of PV. Previous small studies have examined the incidence and effects of HFE C282Y mutations in PV and have not observed associations^55, 56^. However, our study was able to take advantage of the large UK Biobank dataset and utilise an unbiased approach to establish the role of HFE in PV diagnosis.

Given the role of HFE in hepcidin regulation, we dissected the role of hepcidin in PV using novel pre-clinical models. Genetic deletion of hepcidin in our PV model increased erythroid parameters, indicating that in PV, unrestricted erythroid iron access results in unfettered increased hematocrit and hemoglobin production. Conversely, physiologic upregulation of hepcidin expression through RNA interference of *Tmprss6* induced reductions in serum iron, depriving the BM of iron and causing reduced hematocrit, hemoglobin concentration. Hepcidin is thus critical in determining the erythroid phenotype in PV.

The dependence of the PV erythroid phenotype on systemic iron homeostasis provides the mechanistic rationale for the suite of emerging treatments that target iron metabolism for PV. The mainstay of current treatment for PV is venesection to decrease hematocrit below 45%, at which serious thrombotic events become less likely^7, 9^. Venesection lowers the hematocrit by removing red cells and inducing systemic iron deficiency,^19^ thus ameliorating further erythrocyte production^7^. However, venesection can cause unwanted adverse effects including vasovagal reactions due to fluid shifts^57^, as well as symptoms such as chronic fatigue due to systemic iron deficiency^58^, and incurs direct and indirect health care costs associated with patient visits. Some patients are unable to tolerate venesection due to the severity of adverse events or do not achieve satisfactory hematocrit responses and hence require second-line non-targeted cytoreductive agents. New therapeutic options for PV are thus needed. Withholding iron from the BM by inhibiting iron export to the plasma offers an exciting therapeutic opportunity to replace therapeutic venesection to treat PV. We demonstrated that upregulation of endogenous hepcidin levels through liver-specific *TMPRSS6* siRNA deprives the serum of iron and reduces hematocrit and hemoglobin concentration in our PV model. A similar approach using anti-sense oligonucleotide therapies has likewise recently shown promising preclinical results^22^. A clinical candidate of the *TMPRSS6* siRNA used here (SLN124) has been granted Orphan Disease Designation status by the FDA for treatment of PV. A Phase 2 clinical trial showed that a hepcidin mimetic can obviate the need for venesection in previously venesection-dependent PV patients^21^. Our data indicate that liver iron stores were not altered by *TMPRSS6* siRNA treatment, consistent with this therapy redistributing iron stores rather than inducing systemic iron depletion, a feature that could protect patients from symptomatic iron deficiency. *TMPRSS6* siRNA treatment does not affect fluid levels and therefore is unlikely to result in adverse vasovagal responses. Since hepcidin mimetics and *TMPRSS6* siRNA therapies can be administered by subcutaneous injection, these therapies could potentially be self-administered by patients, reducing healthcare costs.

Clonal erythroid expansion in PV results in expansion of later-stage erythroblasts. ERFE is the erythroid-derived negative regulator of hepatic hepcidin expression^15^. Although we detected elevated ERFE in our PV model and in PV patients as would be expected,^59^ *Erfe* deletion did not modify hepcidin expression or alter the disease phenotype in our PV model. These findings may reflect the lesser degree of ERFE elevation in PV compared with diseases of ineffective erythropoiesis such as thalassaemia^60^. Elevations of ERFE in our model of PV were comparable to elevations detected in experimental ‘low’ level ERFE-overexpressing mice, where modest elevations in serum and liver iron were detected, but expression of hepcidin was unchanged^61^.

We hypothesised that in PV, where patients develop iron deficiency and have increased erythropoietic drive, hepcidin would be suppressed. However, this was not the case in our PV mouse model, a finding that recapitulates clinical studies that demonstrate hepcidin levels in newly diagnosed PV patients are comparable to healthy controls^62, 63^. Our data suggest that in PV, hepcidin may be upregulated by inflammation. Numerous inflammatory cytokines (including IL6) are elevated in PV patients, some of which may portend inferior prognosis^48, 50, 64^. Surprisingly, we found IL6 was not the sole driver of hepcidin expression in PV; rather, other IL-6 family cytokines, which signal via GP130 coupled receptors^65^, are implicated. Further characterisation of the role of these cytokine(s) in hepcidin regulation in PV will be an important continuation of this work.

In summary, our findings implicate systemic iron regulation as a key determinant of the clinical severity of PV and lay the foundation for therapeutic strategies that modify iron regulation as potential therapeutics for this disease.

## Acknowledgements

The authors thank the WEHI Bioservices facility for husbandry of all animals and assistance with animal experiments, Stephen Wilcox for assistance with sequencing of RNA indexed libraries, and the WEHI Business Development Office and Legal and Licencing Office for support with Material Transfer Agreements. We thank Prof Hal Drakesmith and Dr Andrew Armitage for sharing iHamp and *Erfe* knockout mice. We thank all the participants who provided samples for this study, and Kelli Grey, Naomi Sprigg and Fiona Sultana for collecting patient samples. The authors would also like to thank our PV patient consumers, Nathalie Cook and Anna Steiner, for their input and feedback on this work.

This work was supported by the National Health and Medical Research Council (GNT1158696, GNT1159171, GNT1195236, GNT2009047, GNT1058344 and GNT1113577). The contents of this published material are solely the responsibility of the individual authors and do not reflect the views of the NHMRC or funding partners. This work was also made possible through the Victorian State Government Operational Infrastructure Support and Australian Government National Health and Medical Research Council (NHMRC) Independent Research Institute Infrastructure Support Scheme (IRIISS). This research has been conducted using the UK Biobank Resource under Application Number 36610. We want to acknowledge the participants and investigators of the FinnGen study.

## Author Contributions

Conceptualized study – C.B., K.B., A.P.N. and S-R.P.; Designed research – C.B., V.E.J., U.S., A.L.G., W.S.A., M.B., A.P.N., and S-R.P.; Performed research – C.B., V.E.J., A.P., T.H., G.M-M., K.F., R.A., D.C. and A.B.; Analyzed data – C.B., V.E.J., A.L.G and C.S.N.L-W-S; Wrote the manuscript - C.B., V.E.J., U.S., R.A., D.C., A.B., W.S.A, M.B., A.P.N. and S-R.P.

## Conflict of Interest

U.S. is a full-time employee of Silence Therapeutics GmbH.

