## Supplemental Material for "Iron homeostasis governs erythroid phenotype in Polycythemia Vera"

### Supplemental Methods

*UK Biobank – defining Polycythemia Vera cases, and blood cell trait phenotypes*

Data from British individuals in the UK Biobank dataset[1] were analysed. PV cases were identified using Hospital Episode Statistics (HES). Individuals were classified as a case if they had a relevant ICD code (ICD-10 of D45 or an ICD-9 of 238.4) recorded under any of the following: Underlying (primary) cause of death; contributory (secondary) causes of death; Diagnoses (main); Diagnoses (secondary). Control individuals who did not have an ICD-9 or -10 code relating to any Myeloproliferative Neoplasm (PV - as above, Essential Thrombocythemia ICD-10, D47.3/ ICD-9, 238.71 or Primary Myelofibrosis ICD-10, D47.4 / ICD-9, 289.83). Individuals without HES data available were excluded.

Cleaning of four blood cell traits (hemoglobin concentration, hematocrit percentage, mean corpuscular volume, and erythrocyte count) was undertaken as previously reported[2]. Briefly, all traits were first log-transformed, or logit transformed (hematocrit percentage). Using a restricted set of central measurements (measures <3.5 median absolute deviations from the median), a generalised additive model (GAM, using the mgcv R package) was fitted, to model the effects of several technical covariates (time of measurement, instrument, acquisition route, day of the week). In the full dataset, model residuals were calculated and used to generate traits adjusted for technical effects. Finally, outliers (>6 median absolute deviations from the median) were excluded. The log (or logit) transformed, adjusted traits were used for downstream genetic analyses.

*UK Biobank – genetic data quality control and genome-wide association analysis*

The genotyping procedure, quality control and imputation of the UK Biobank cohort is described in detail elsewhere[1]. Individuals with outlying heterozygosity or high levels of missingness were excluded prior to imputation. We further excluded individuals who had withdrawn consent, samples where the self-reported sex did not match the genetically inferred sex, samples with putative sex chromosome aneuploidy, and samples with an excess of relatives (>10 3rd degree). Samples were then restricted to the "White British" subset[1]. Variants were filtered to include those with MAF>0.01%, and INFO>0.8.

Source code can be accessed at: https://github.com/bahlolab/polycythemiaVeraGWAS

*Generation of Cre recombinase-inducible Jak2-V617F transgenic mice.*

Complementary DNA (cDNA) encoding murine *Jak2*-V617F was cloned downstream of a CMV enhancer-chicken beta-actin promoter element with an intervening cassette including 3 transcriptional termination elements flanked by loxP sites (modification of plasmid pCAGGS-loxSTOPlox-ClaI flp-in, obtained from Addgene; deposited by Rudolf Jaenisch). This was then inserted downstream of the *Col1a1* locus by FRT/Flpe recombinase-mediated site-specific integration by co-transfection with a Flpe expression vector into C57BL/6 5B3 ES cells containing an FRT-hygro-pA "homing" cassette at the ColA1 gene. Correctly targeted ES cell clones were injected into Balb/c blastocysts to generate chimeric mice. Male chimeras were mated with C57BL/6 females to yield heterozygotes for the targeted allele, referred to as LSL-Jak2-V617F mice. To enable tamoxifen-dependent activation of Jak2-V617F expression, LSL-Jak2-V617F mice were intercrossed with previously described CreERT2 mice[3].

*Bone marrow transplant model of Polycythemia Vera*

Bone marrow cells were flushed from the tibias and femurs of LSL-Jak2-V617F; $CreERT2^{T/+}$ mice (or LSL-Jak2-V617F lacking CreERT2 for controls) with 5% FBS (Gibco) in KDS BSS (150mM Sodium Chloride, 3.7mM Potassium Chloride, 2.5mM Calcium Chloride Dihydrate, 1.2mM Magnesium Sulfate Heptahydrate, 14.8mM HEPES) and passed through a 23G needle to produce a single cell solution under sterile conditions. Bone marrow cells were then pelleted under centrifugation (470g, 5min) and resuspended in 2% FBS/KDS BSS. $2.5x10^6$ donor bone marrow cells were intravenously injected into lethally irradiated (two 5.5 Gy doses given 4 hours apart) Ly5.1/J (B6.SJL-Ptprca Pepcb/BoyJ) recipient mice. Irradiated mice were given antibiotics (1.1mg/ml neomycin) in their drinking water for 3 weeks post irradiation. Seven weeks post bone marrow transplantation, mice were given tamoxifen (Sigma; 4.2mg in 90% corn oil/10% ethanol) by oral gavage on 2 consecutive days to induce expression of the mutant *Jak2* allele. Mice were then humanely euthanised 8-10 weeks later.

*Chimerism assessment*

Six and a half weeks after bone marrow transplantation, mice were bled from the retro-orbital plexus into Microvette® 500 K3 EDTA tubes (Sarstedt). 100µl blood was added to 10ml red cell lysis buffer (156mM Ammonium Chloride, 11.9mM Sodium Bicarbonate, 0.097mM EDTA) and then pelleted under centrifugation (470g, 5min). The cell pellet was washed once in 4.5ml FACS buffer (2% FBS, 0.002% Sodium Azide in KDB BSS), then resuspended in FACS buffer containing CD45.1-PE (clone A20, made in house) and CD45.2-A647 (clone S450-15.2, made in house) and incubated on ice for 30 mins. Cells were washed once in FACS buffer and resuspended in 100µl FACS buffer and 50uL Hydroxystilbamidine (25µg/mL; Biotium). Samples were run on a BD FACSymphony A3 using BD Diva software. Samples were analysed using FlowJo 10.8.0 (BD). Chimerism was defined as the percentage of live cells expressing CD45.2.

*Gene expression analysis*

Total RNA from liver, spleen, kidney, and bone marrow samples and HepG2 and Huh7 cells were isolated using the ISOLATE II RNA Mini Kit (Bioline). 500ng RNA was reverse-transcribed to cDNA using the SensiFAST cDNA synthesis kit (Bioline), according to the manufacturer's instructions. Gene expression levels were measured by RT-qPCR using SensiFAST SYBR No-ROX kit or SensiFAST Probe No-ROX kit (both Bioline) on a LightCycler® 480 II (Roche). Primers or probes used for RT-qPCR are found in Supplemental Table 5.

*Analysis of bone marrow erythropoiesis by FACS*

Bone marrow cells were flushed from the tibias and femurs and a single cell solution created as above. $2x10^6$ cells were pelleted under centrifugation (470g, 5min, $4^oC$), resuspended in 50µl FACS buffer containing anti-mouse CD44-FITC (clone: IM781, made in house) and anti-mouse TER119-APC (clone TER119, made in house) and incubated on ice in the dark for 30 minutes. Cells were washed in 4.5ml FACS buffer, pelleted under centrifugation (470g, 5min, $4^oC$) and resuspended in 100µL FACS buffer. 50µL Hydroxystilbamidine (25µg/mL – Biotium) was then added before running samples on a BD FACSymphony cytometer using BD Diva software. Sample analysis was performed using FlowJo 10.8.0 (BD). Gating of erythroid cell populations has been previously described[4].

*RNA-Seq*

300ng RNA was used as input for purifying the poly(A)-containing mRNA molecules, RNA amplification, and synthesis of double-stranded cDNAs according to Illumina's TruSeq RNA Sample Prep guidelines. The indexed libraries were pooled and diluted to 1.5pM for paired end sequencing (2x 76 cycles) on a NextSeq 500 instrument using the v2 150 cycle High Output kit (Illumina) as per manufacturer's instructions. The base calling and quality scoring were determined using Real-Time Analysis on board software v2.4.6, while the FASTQ file generation and de-multiplexing utilised bcl2fastq conversion software v2.15.0.4.

Paired-end RNA sequencing reads were aligned to the mm10 build of the mouse reference genome using the Rsubread package v2.4.3[5]. Over 98% of fragments (read pairs) mapped to the reference genome for each sample. Successfully mapped fragments were then summarized into gene-level counts using featureCounts[6] and genes were identified using Gencode annotation to the mm10 genome (version M25)[7] where 83-86% of mapped fragments were assigned to genes. Differential expression analyses were carried out using limma v3.48.3[8,9] and edgeR v3.34.0[10].

After excluding genes that were to be experimentally confirmed, expression-based filtering was carried out using the filterByExpr function with default settings in edgeR. A total of 15,690 genes remained for downstream analysis. Compositional differences between the libraries were normalized using the trimmed mean of M-values (TMM) method[11]. Counts were then transformed to $\log_2$ counts per million (logCPM) with associated observational-level weights using voom[12]. Differential expression between PV and control samples was assessed using linear models and robust empirical Bayes moderated t-statistics. Furthermore, the linear models incorporated a batch effect correction for assay effect to increase precision. False discovery rate was controlled below 5% using the Benjamini and Hochberg method. The mean-difference plot was generated using the limma's plotMD function while the heatmap was created using the pheatmap software package.

*Analysis of hematological and iron parameters*

Full blood counts were determined using an Advia2120i on EDTA anticoagulated blood. Serum iron was measured using an Abbott ARCHITECT analyser using the MULTIGENT Iron assay. Tissue non-heme iron was measured as previously described[13]. Hepcidin (Intrinsic Life Sciences), erythroferrone (Intrinsic Life Sciences) and interleukin-6 (R&D Systems) ELISAs were performed following manufacturer's instructions.

*Cell culture*

HepG2 and Huh7 cells were maintained in Dulbecco's modified Eagle medium (DMEM; Gibco) containing 1g/L D-Glucose, 1% L-glutamine, 100mg/L Sodium Pyruvate and supplemented with 1% penicillin-streptomycin and 10% FBS (Gibco). $3x10^5$ cells were seeded in wells of 24-well tissue culture treated plates. After 24h, 10ng/ml recombinant human IL6-family cytokines (IL6, IL11, OSM, LIF, IL27, CT-1, CTNF and CLCF1 all R&D Systems) were added to the media; or media was removed and replaced with media supplemented with 2% human plasma (instead of FBS) with or without the addition of 300ng/ml anti-human GP130 (clone 28105; R&D Systems). In all cases cells were lysed 24 hours later and RNA isolated for RT-qPCR as above.

### Supplemental Tables

Supplemental Tables 1 to 5 are supplied as an addition excel file

### Supplemental Figures

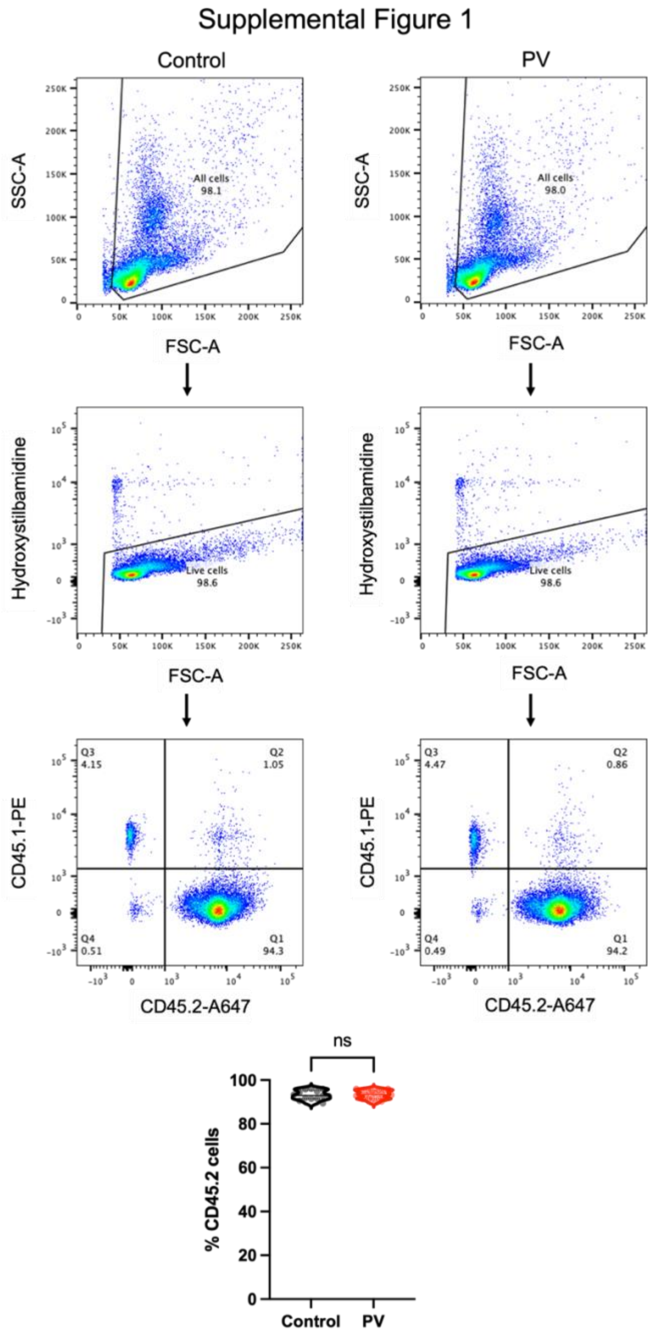


**Supplemental Figure 1 – Bone marrow transplanted mice are highly chimeric.**

Chimerism, defined as percentage of CD45.2 live nucleated blood cells, of control and PV mice determined by flow assisted cell sorting of peripheral blood cells. Top plot, gating of non-fragmented cells. Middle plot, gating of live (Hydroxystilbamidine negative) cells. Bottom plot, expression of CD45.1 and CD45.2 of live cells. N=13 control/10 PV. Unpaired 2-tailed t-test with Welch's correction. ns= non-significant.
